# Activator-binding domains are required but insufficient for Med15 recruitment to UAS, revealing a critical role for its disordered C-terminus in directing genome targeting

**DOI:** 10.64898/2026.05.24.727475

**Authors:** Aileen Cohen, Joshua Bugis, Vladimir Mindel, Gilad Yaakov, Naama Barkai

## Abstract

The Mediator complex is a conserved transcriptional coactivator that bridges transcription factors (TFs) with the core transcriptional machinery. In budding yeast, TFs recruit the Mediator to upstream activation sequences (UASs) by interacting with three activator-binding domains (ABDs) within the Med15 tail subunit. However, these ABD-TF interactions exhibit relatively low affinity, raising the question of whether they fully account for Med15 recruitment in vivo. Here, we address this question by combining genome-wide profiling with systematic mutational analysis. We demonstrate that while the three ABDs are collectively required for Med15 recruitment, they are not individually required and are not sufficient to specify the genomic binding pattern of Med15. Instead, removal of an intrinsically disordered C-terminal redistributes Med15 occupancy across UASs. Notably, C-terminal truncation alters Med15 binding even at TF-depleted regions, and these effects are recapitulated by deleting Med2 or Med3, which further stabilize TF-dependent recruitment. Together, these findings support a dual mechanism of Med15 UAS recruitment, whereby TF-ABD recruitment is directed and stabilized by C-terminal mediated interactions.

## Introduction

Transcription factors (TFs) orchestrate gene expression by binding specific cis-regulatory regions within the genome. Upon binding, TFs recruit general coactivators to their target sites, facilitating assembly of the pre-initiation complex (PIC) and initiating transcription^1,2^. A central coactivator conserved from yeast to humans is the Mediator complex^3–6^. This large, multi-subunit complex is organized into a core and a tail module. Core Mediator is broadly required for transcription and can associate with gene promoters, located at the transcription start site (TSS), independently of the tail module. In contrast, the tail module is required for TF-dependent recruitment of Mediator to upstream activating sequences (UASs) located few hundred base-pairs (bps) away from the TSS, and for the consequential induced gene activation. Hence, interactions between UAS-bound TFs and the Mediator tail constitute a critical step in gene induction. In budding yeast, the Med15 subunit functions as a major TF interaction hub, mediating Mediator recruitment to UAS regions^7^.

Med15 interacts with TFs primarily through its N-terminal domain, whereas its C-terminal region is required for transcriptional activation^8,9^ and for maintaining Mediator complex integrity^8,10^. Within the N-terminus of Med15, three activation-binding domains (ABDs) that engage the activation domains (ADs) of multiple TFs^10–17^ have been identified. However, the affinities of these ABD-AD interactions are typically in the micromolar range^14,15,17–19^, in contrast to the nanomolar affinities commonly observed for TF-DNA and TF-repressor interactions (e.g., Gal4-Gal80 ^20^). This discrepancy raises the question of whether the relatively weak ABD-TF interactions are sufficient to account for Med15 recruitment to TF-bound UAS elements *in vivo*, or whether additional interactions contribute to Med15 targeting across the genome.

In this work, we systematically searched for determinants of Med15 recruitment to UASs, refining the contributions of the three ABDs and testing whether additional elements contribute. We find substantial redundancy among the ABDs and show that, while collectively required, they are not sufficient to explain Med15 targeting but act in cooperation with a disordered C-terminal region that shapes Med15 binding across UAS. Together, these results point to a more complex logic for Med15 targeting in vivo that goes beyond simple TF-mediated recruitment.

## Materials and methods

### Design of strains used in the study

Genetic manipulations were performed on the S. cerevisiae BY4741 strain (genotype: MATa his3Δ1 leu2Δ0 met15Δ0 ura3Δ0 genotype) using CRISPR with a genotype based on the sequence cofactor-MNase-SV40NLS-Adh1 terminator. All strains generated for this study were verified using PCR and gel electrophoresis followed by Sanger DNA sequencing. Supplementary Table S1 contains information on all strains used in the study.

### Budding yeast growth, maintenance, and genetic manipulation

Transformations were performed using the LiAc/SS DNA/PEG method. In strains deleted of the 12 stress TFs, the transformation mix was supplemented with DMSO in a final concentration of 5% prior to the heat shock. Following validation by Sanger sequencing, the pbRA89 (Addgene plasmid #100950) carrying the CRISPR-Cas9 system from positive colonies were lost by growth in YPD media (yeast extract, peptone, dextrose) overnight, and selection for colonies without bRA89-encoded Hygromycin resistance. Ligation of the gene-specific guide-RNA into the bRA89 plasmid was performed as previously described.

### Cell growth before experiments

Yeast strains were freshly thawed from frozen stock, plated on YPD plates, and grown until visually distinct colonies were seen on the plate. Single colonies were picked and grown at 30 °C in liquid SD medium (synthetic complete with dextrose) overnight, reaching the stationary phase (OD600 ≈10), then diluted again into fresh SD medium for the experiment.

### ChEC-seq experiments

The experiments were performed as described previously (see ref 24), with some modifications. The stationary cultures described above were diluted ~2 × 10^3^-fold into 5 ml fresh SD media and grown overnight to reach an OD600 of 4 the following morning. Cultures were pelleted at 4000g for 2 min and resuspended in 0.5 ml buffer A (15 mM Tris pH 7.5, 80 mM KCl, 0.1 mM EGTA, 0.2 mM spermine, 0.5 mM spermidine, 1× cOmplete EDTA-free protease inhibitors [Roche, one tablet per 50 ml buffer], 1 mM PMSF) and then transferred to 2 ml 96-well plates (Thermo Scientific). Cells were washed twice in 1 ml Buffer A. Next, the cells were resuspended in 150 μl Buffer A containing 0.1% digitonin, transferred to an Eppendorf 96-well plate (Eppendorf), and incubated at 30°C for 5 min for permeabilization. Next, CaCl2 was added to a final concentration of 2 mM to activate the MNase and incubated for 30 seconds exactly. The MNase activation was stopped by adding an equal volume of stop buffer (400 mM NaCl, 20 mM EDTA, 4 mM EGTA, and 1% SDS) to the cell suspension. After this, the cells were treated with Proteinase K (0.5 mg/ml) at 55°C for 30 min. An equal volume of phenol-chloroform pH 8 (Sigma-Aldrich) was added, vigorously vortexed, and centrifuged at 17000g for 15 minutes to extract DNA. Following the extraction, the DNA was precipitated with 3 volumes of cold 96% EtOH, 45 mg Glycoblue, and 20 mM sodium acetate at –80°C for >1 h. Next, the tubes were centrifuged (17000g, 4°C for 10 min), the supernatant was removed, and the DNA pellet was washed with 70% EtOH. The DNA pellets were dried and resuspended in 30 μl RNase A solution (0.33 mg/ml RNase A in Tris-EDTA [TE] buffer [10 mM Tris and 1 mM EDTA]) and incubated at 37°C for 20 min. DNA cleanup was performed using SPRI beads (Ampure XP, Beckman Coulter) to enrich small DNA fragments and remove large DNA fragments that might result from spontaneous DNA breaks. First, a reverse SPRI cleanup was performed by adding 0.8× (24 μl) SPRI beads, followed by 5 min incubation at RT. The supernatant was collected, and the remaining small DNA fragments were purified by adding additional 1× (30 μl) SPRI beads and 5.4× (162 μl) isopropanol and incubating 5 min at RT. The beads were washed twice with 85% EtOH, and DNA was eluted in 30 μl of 0.1× TE buffer.

### ChEC-Seq next-generation sequencing library preparation

Library preparation was performed as described in 28. Following RNase treatment and reverse SPRI cleanup, the DNA fragments served as an input to an end-repair and A-tailing (ERA) reaction. 5.4 μl ERA reaction was prepared (1 × T4 DNA ligase buffer [NEB], 0.5 mM dNTPs, 0.25 mM ATP, 2.75% PEG 4000, 6U T4 PNK [NEB], 0.5U T4 DNA Polymerase [Thermo Scientific], 0.5U Taq DNA polymerase [Bioline]) and added to 14.6 μl of each sample and incubated for 20 min at 12°C, 15 min at 37°C and 45 min at 58°C in a thermocycler. After the ERA reaction, reverse SPRI cleanup was performed by adding 0.5× (10 μl) SPRI beads (Ampure XP, Beckman Coulter). The supernatant was collected, and the remaining small DNA fragments were purified with an additional 1.3× (26 μl) SPRI beads and 5.4× (108 μl) isopropanol. After washing with 85% EtOH, small fragments were eluted in 17 μl of 0.1 × TE buffer; 16.4 μl elution was taken into 40 μl ligation reaction (1 × Quick ligase buffer [NEB], 4000U Quick ligase [NEB], and 6.4 nM Y-shaped barcode adaptors with T-overhang (52) and incubated for 15 min at 20°C in a thermocycler. After incubation, the ligation reaction was cleaned by performing a double SPRI cleanup: first, a regular 1.2× (48 μl) SPRI cleanup was performed and eluted in 30 μl 0.1 × TE buffer. Then instead of separating the beads, an additional SPRI cleanup was performed by adding 1.3× (39 μl) HXN buffer (2.5 M NaCl, 20% PEG 8000) and final elution in 24 μl 0.1 × TE buffer; 23 μl elution were taken into 50 μl enrichment PCR reaction (1 × Kappa HIFI [Roche], 0.32 μM barcoded Fwd primer and 0.32 μM barcoded Rev primer 29, and incubated for 45 s in 98°C, 16 cycles of 15 s in 98°C and 15 s in 60°C, and a final elongation step of 1 min at 72°C in a thermocycler.

The final libraries were cleaned using 1.1× (55 μl) SPRI and eluted in 15 μl 0.1 × TE buffer. Library concentration and size distribution were quantified by Qbit (Thermo Scientific) and TapeStation (Agilent). For multiplexed next-generation sequencing (NGS), all barcoded libraries were pooled in equal amounts, and the final pool was diluted to 2 nM and sequenced on NovaSeq 6000 (Illumina)/ NovaSeq X (Illumina). Sequence parameters were Read1: 61 nucleotides (nt), Index1: 8 nt, Index2: 8 nt, Read2: 61 nt.

### ChEC-seq NGS data processing

Raw reads from ChEC-seq libraries were demultiplexed using bcl2fastq (Illumina), and adaptor dimers and short reads were filtered out using cutadapt with parameters: ‘−O 10 –pair-filter = any –max-n 0.8 –action = mask’. Filtered reads were subsequently aligned to the S. cerevisiae genome R64-1-1 using Bowtie 2 with the options ‘--end-to-end --trim-to 40 --very-sensitive’. The genome coverage of fully aligned read pairs was calculated with GenomeCoverage from BEDTools using the parameters ‘-d –5 –fs 1’, resulting in the position of the fragment ends, which correspond to the actual MNase cutting sites. The reads were normalized to 10^7^.

### Promoter definition

Promoters were defined only for genes with an annotated transcript, as described before. The TSS was defined using these parameters (see ref 20). The length of each promoter was defined as 700 bp upstream of the transcription start site (TSS) or to the position where a promoter meets another transcript. The signal across each promoter was summed and normalized to the maximal promoter length 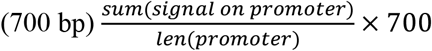 to calculate the overall promoter binding for each sample.

### Target promoter definition

To define the target promoters of analyzed strains (Figures 1B-E, 2A,2C, 3C-E, 4B-F, S1B, S2B-C), we calculated the sum of signal on defined promoters (see Promoter definition), Z-scores were calculated, and promoters with a Z-score ≥3 were selected as the respective targets of Med15 in WT, Med15-ΔC400, and in Δ12 TFs Med15-DBD_Msn2_ in Δ12 TFs. For the lowest targets, Z-score data frames were sorted, and the intersection between the lowest targets and the top targets (already calculated) was checked (e.g, the lowest in Med15 Δ12 TFs was intersected with top Med15 in WT and with top in Med15-DBD_Msn2_ to define the targets lost and restored).

**Figure 1:**
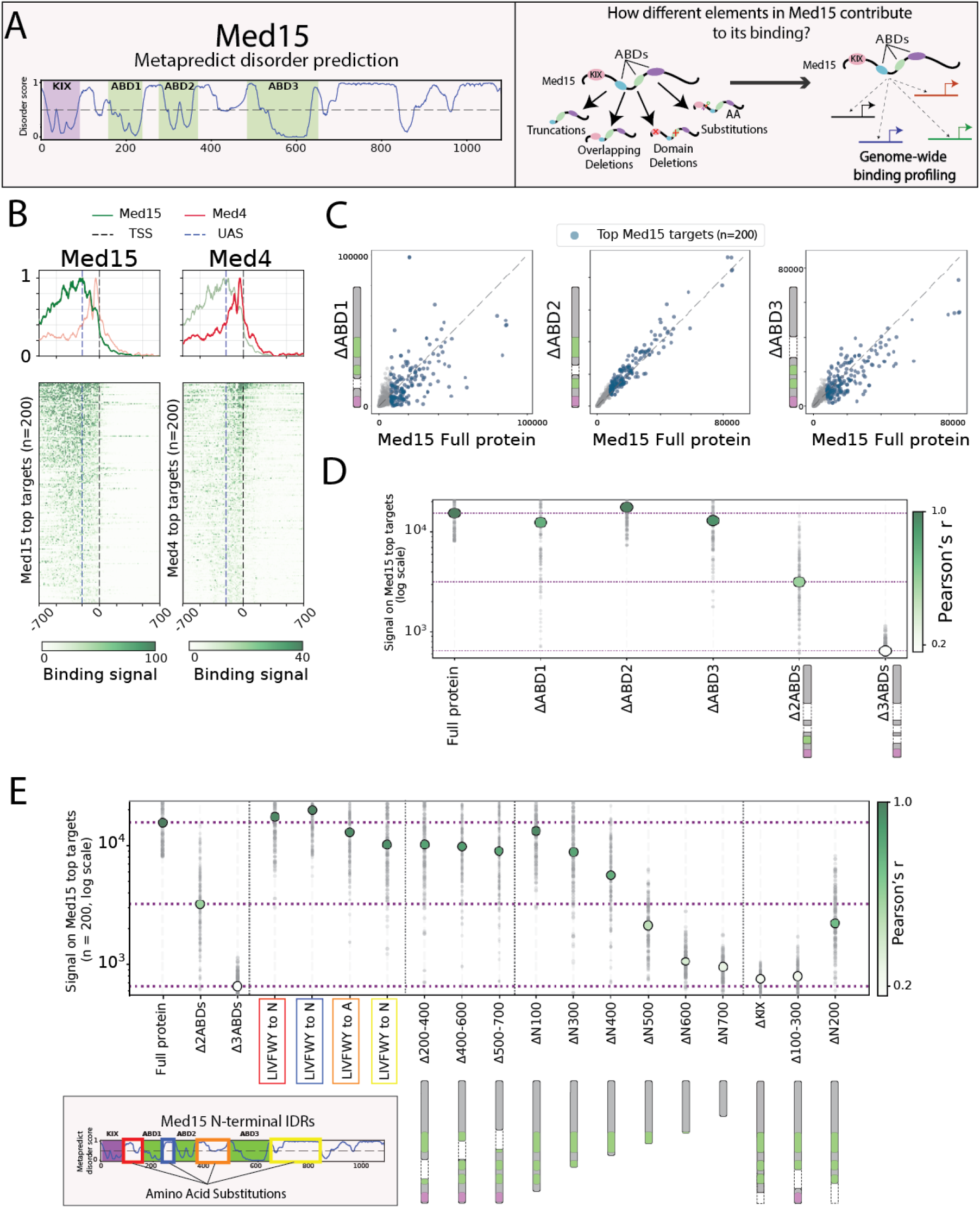
The ABDs in the N-terminal domain act redundantly. (A) *Schemes*. (left) Med15 has multiple regions that are predicted to be disordered using Metapredict (Version 2.63) and a few studied domains (KIX, ABDs) predicted to be ordered. (right) The experimental question and design. We tested the genome-wide binding of Med15 as a full protein and with various mutations such as truncations (from the N or C-terminus), domain and region deletions, and amino acid substitutions. (B) *Med15 and Med4 bind on different locations in regulatory regions*. Binding signals of Med15 and Med4 (green and red lines, respectively) at their top 200 targets, are shown as heatmaps and meta profiles centered around the TSS and flanked by 700bp regions upstream and downstream (methods, Table S1). Color bars represent signal strength on each bp in the sequence. (C) *Deleting individual ABDs has a limited effect on the binding pattern of Med15*. Promoter binding signals of the indicated strains across all yeast promoters are shown. The diagonal dashed line indicates x=y. Blue dots represent the top Med15 targets shown in (B). Schemes of the deleted ABD correspond to the structure of Med15 shown in (A). (D) *Deleting two ABDs reduces Med15’s binding to its top targets, and deleting all three ABDs abolishes binding*. Median signal (circles, in log scale) at Med15 top targets in the indicated strains. The color intensity corresponds to Pearson’s correlation with the full Med15. The grey dots represent the spread of binding on the indicated targets shown in (B). The purple horizontal lines mark the binding of the full protein, ΔABD2 ΔABD3 (abbreviated as Δ2ABDs) and ΔABD1 ΔABD2 ΔABD3 (abbreviated as Δ3ABDs). (E) *Med15 recruitment across the genome is dependent on the ABDs and their flanking IDRs*. Shown is the median signal on Med15’s top targets in the indicated strains, including domain deletions (e.g, ABD deletions), serial truncations, 200AA region deletions, and amino acid substitutions (in log scale) as in (D). (bottom) N-terminal disordered regions that were mutated using AA substitutions, each represented by a different corresponding color.

### Binding around the TSS

The ChEC-seq signal on target promoters of tested strains (Fig. 1B) was extracted ±700bp surrounding the respective TSS. The signal was calculated and was then averaged by a rolling window of 10 and plotted as a heatmap (see Target promoter definition).

### Averaged signal on target promoters

The ChEC-seq signal on target promoters of all tested mutations across the yeast genome was extracted and then averaged by a rolling window of 30 and was plotted in 1400 bases centered around the TSS of target promoters, the signals were normalized to the peak of each of the strains to allow comparison (Figures 1B, 2A, 3B, S3, see Target promoter definition).

### Flocculation assay

Yeast strains were grown overnight at 30 °C in YPD/SD with shaking until they reached stationary phase. Cultures were then diluted in fresh YPD/SD (according to initial growth media) to OD 0.1 in the morning and grew to OD 0.4-0.6. Then the cells were diluted in the respective media at a 1:10 ratio and transferred to a cuvette. Optical density (OD) at 600nm was measured every minute for 30 minutes using a spectrophotometer. A decrease in OD over time was used as an indicator of flocculation, with more rapid decreases corresponding to stronger flocculation phenotypes that lead to faster settling of the undisturbed cells (Fig. S1D).

## Results

### The three Med15 activator-binding domains (ABDs) act redundantly in genome targeting

The relatively low affinity of Med15 ABDs for TF activation domains (ADs) prompted us to investigate whether these ABDs are sufficient to account for Med15 recruitment to TF-bound upstream activation sequences (UASs). To address this, we employed genome-wide profiling as our primary approach. Previous studies have demonstrated that Mediator localizes to both gene promoters and UASs via two distinct mechanisms. While promoter binding is independent of the tail module, it is highly transient; consequently, it is undetectable when profiling Med15 in wild-type cells and requires specific molecular perturbations to be visualized. In contrast, Mediator recruitment to UASs is mediated by its tail module, resulting in a more stable interaction that is readily observed during Med15 profiling in wild-type background^21–26^.

We mapped Med15 binding across the genome using ChEC-seq (Fig. 1A). As we showed previously^25,26^, the Med15 ChEC-seq profile is consistent with ChIP-seq mapping and captures its recruitment to TF-bound UASs at tail-dependent genes^21–23,27^. Note that, operationally, we define promoters as the regions immediately flanking the transcription start site (TSS) present across all genes, whereas UASs refer to regions harboring binding sites for Med15-recruiting TFs, which are typically located 200–400 bp upstream of the TSS and are found at only a subset of genes. Indeed, Med15 binding was preferentially located at ~200bp upstream to the TSS, in locations dense with TF-binding sites, contrasting other mediator components that displayed prominent binding at 50 bp of the TSS (Fig. 1B, Fig. S1A).

Given the relatively weak ABD-AD interactions, we expected the three ABDs to cooperate and be essential in Med15 recruitment. However, removal of each individual ABDs, revealed that none was required. First, we noted that, in all mutants, binding peaks remained at similar ~200 bp distance from the TSS, consistent with UAS binding. Second, to examine changes in preferences, we compared the overall binding signals associated with each gene (taking the 700bp upstream of each TSS). This revealed that deletion of ABD1 reduced Med15 binding at a subset of gene targets, whereas deletion of either ABD2 or ABD3 had no or a moderate effect on binding across all its targets (Fig. 1C). Further, ABD2 was of no effect also when deleted in ABD1-mutated cells (FigS.1B). Only upon co-deletion of ABD2 and ABD3, we observed a marked reduction in Med15 targets, and the residual binding was abolished upon removal of all three ABDs (Fig. 1D). Together, these results reveal substantial redundancy amongst the three ABDs in targeting Med15 to its UAS sites.

We further examined the interplay between the three ABDs and their surrounding elements by systematically mutating the 600-residue N-terminus of Med15 where the three ABDs are located (Fig. 1E). First, we noted that Med15 binding was largely invariant to mutations that altered, individually, the ABD-flanking intrinsically disordered regions (IDRs) (Fig. 1E, Fig. S1C). Second, we sequentially removed 100-residue segments from the N-terminus and then deleted overlapping 200-residue segments. Three truncations, including the terminal KIX domain, induced flocculation of the yeast and could not be profiled, but were rescued by longer truncations (Fig. S1D). The remaining truncations progressively abolished Med15 binding across all its targets, with no new targets appearing. Together, these results confirm that the three ABDs are collectively required for Med15 recruitment to its UAS targets while highlighting substantial redundancy amongst them.

### Med15 targeting is controlled by its C-terminus

We next asked whether the N-terminal region containing the three ABDs is sufficient, by itself, for Med15 targeting. For this, we tested a Med15 mutant truncated of its 400 C-terminal amino acids but retaining all three ABDs. This mutant localized reproducibly across the genome (Fig. S1E) and was preferentially associated with UAS regions upstream of the TSS (Fig. 2A) Yet, when examining its binding preferences, we noted that these differed the full-length Med15: the truncation reduced binding at a substantial fraction of target genes and increased at others (Fig. 2A). In addition, the C-terminal appeared to synergize with elements of the N-terminal: extending the C-terminal truncation to include regions that by themselves had little effect abolished Med15 binding altogether (Fig. 2B, C). We conclude that, in the absence of the C-terminus, the three ABDs are not sufficient to specify Med15 binding across the genome.

**Figure 2:**
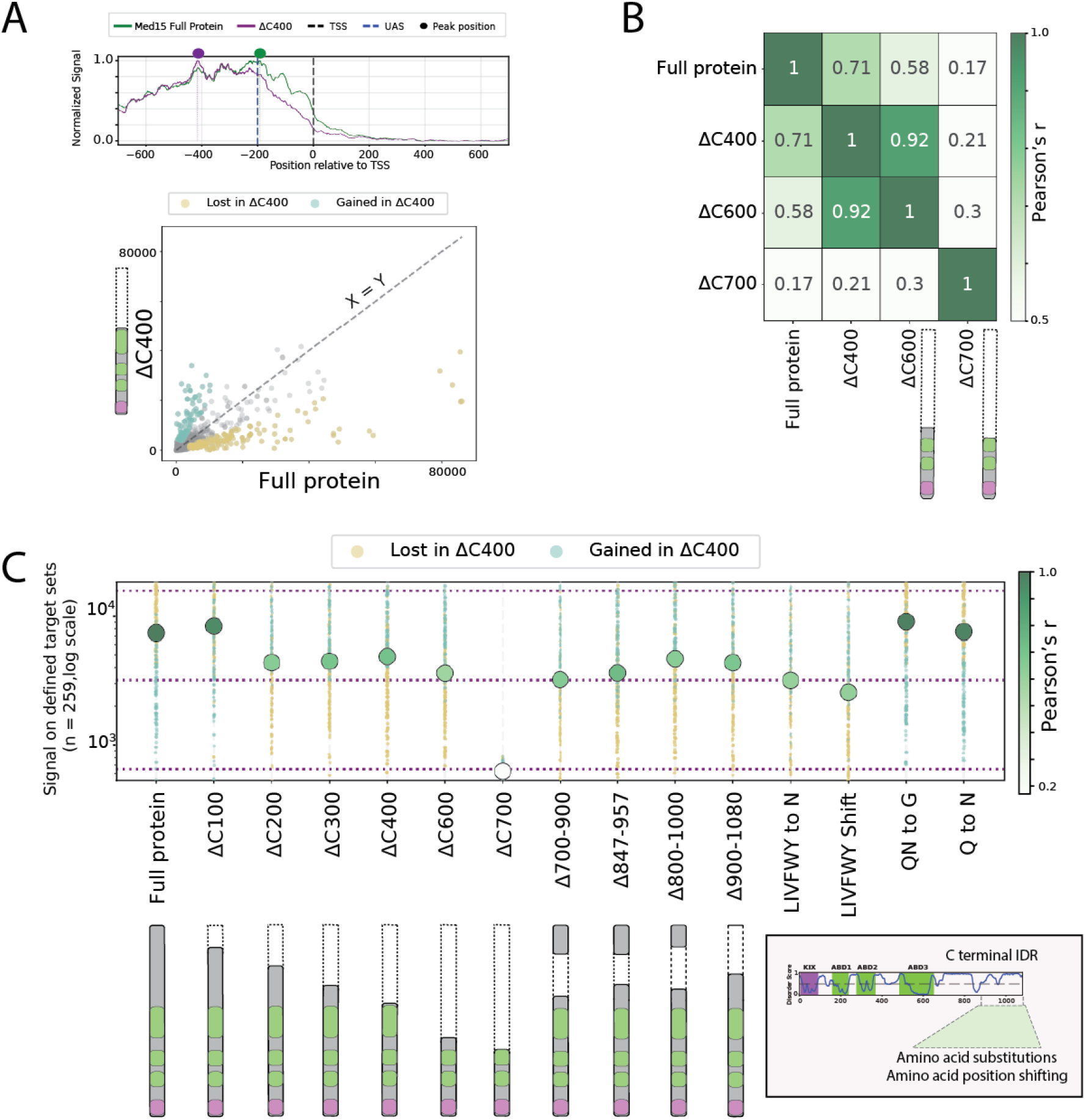
The C-terminus of Med15 contributes to its target selection. (A) *Deleting the C-terminus of Med15 changes its binding preferences and localization*. (top) An averaging of signal on top Med15 targets comparing the binding localization of the full Med15 (green) and the mutated Med15 (purple) on regulatory regions. (bottom) Comparison of the binding signal across all promoters in the indicated strains. The light blue points represent the targets that increased in binding when 400AAs are deleted, while the yellow points represent the targets that were decreased in binding in this mutation (methods, Table S1). Schematic as in Fig. 1C. (B) *Removing ABD3 and its flanking IDRs abolishes binding*. Shown are Pearson’s correlation coefficients comparing the binding signal of corresponding strains. (C) *The hydrophobic residues in the C-terminal region affect Med15’s binding pattern*. Shown is the median signal (in log scale) of indicated mutations on the increased and decreased targets when the C-terminus is deleted (as in (A)). The color intensity corresponds to Pearson’s correlation to the binding of the full Med15 on the represented targets. A scheme of the C-terminal disordered region in which AAs were shifted in locations preserving composition (hydrophobic) or were replaced (hydrophobic by N, QN by G, or Q replaced by N). The representation of the colors and the horizontal lines are as in Fig. 1D.

To refine the functional region within the Med15 C-terminus, we tested a series of smaller truncations. This narrowed the critical region whose deletion still perturbed Med15 binding to a ~100-amino acid segment (residues 800–900) overlapping the MAD (Mediator Activation Domain^28,29^). As this region is predicted to be intrinsically disordered, we tested its sequence requirements by mutating different classes of residue, including charged, hydrophobic or polar. Of these, hydrophobic residues emerged as critical: both the substitution or the rearrangement of these residues recapitulated the phenotype of the full C-terminal deletion, suggesting a role for a short linear motif or a transiently formed structure within the disordered segment ^30^ (Fig. 2C). Together, these results indicate that a disordered region in the Med15 C-terminus complements the ABDs in shaping Med15 targeting.

### The C-terminal binding phenotype may be explained by Med2/3 interactions

We noted that C-terminal truncation shifted Med15 binding further upstream of the TSS, resulting in reduced occupancy at TSS-proximal promoter regions (Fig. 2A). Because this C-terminal region contacts other Mediator subunits^28,29^ and is required for Mediator integrity^7^. This spatial shift could be explained by a weakened interaction with the core Mediator complex. Consistent with this hypothesis, deletion of *MED15* significantly depleted other Mediator subunits from Med15 target sites (Fig. S2A). These observations led us to consider a model wherein the core Mediator stabilizes Med15 binding at a subset of targets through interactions with its C-terminus, explaining the C-terminus contribution to Med15 targeting. To test this model, we deleted *MED16*, a subunit previously implicated in coupling the tail module to the core Mediator^23,31^. However, contrary to our expectation, *med16Δ* cells exhibited little effect on genome-wide Med15 binding, altering neither its target selection nor its preferred positioning relative to the TSS (Fig. 3B). Therefore, coupling to the core Mediator is unlikely to account for the impact of the C-terminus on Med15 recruitment.

**Figure 3:**
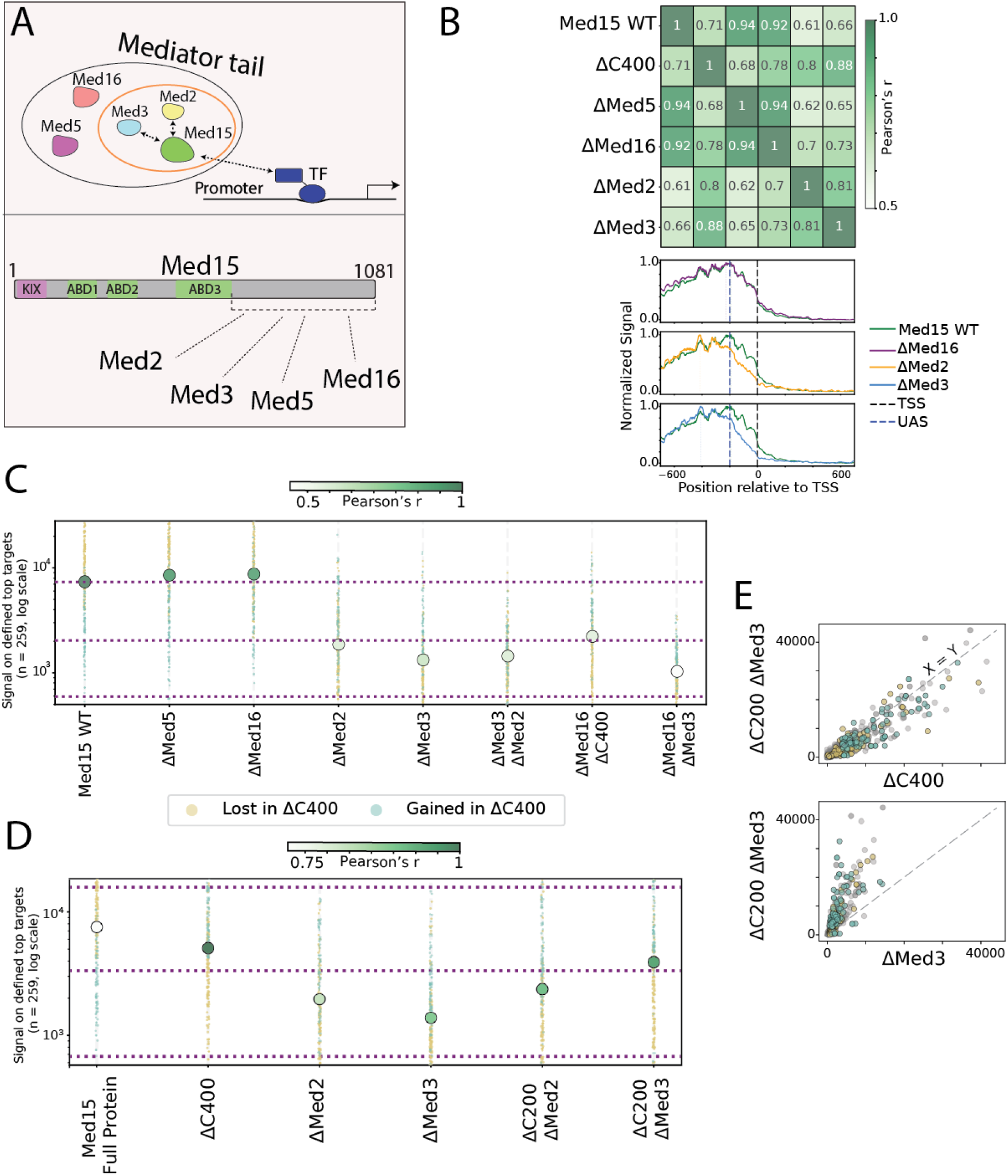
Med15’s target selection depends on its MAD and on the interaction between the MAD, Med2, and Med3. (A) *Med15 communicates with the entire tail module via its C-terminus*. Med15 is a subunit of the mediator tail module. Within the tail, it forms a subcomplex with Med2/3 and further interacts with Med5/16, which connects with other mediator subunits. These interactions occur via the C-terminus of Med15. (B) *The deletion of Med2/3 affects Med15’s binding*. (top) Pearson’s r correlation coefficient comparing the binding signal of Med15 in wild-type versus deletions of tail subunits. (bottom) An averaging of signal on top Med15 targets comparing the binding localization of the indicated mutations on regulatory regions. (C) *Deleting Med2/3 is similar in phenotype to deleting the C-terminus of Med15*. Shown is the median signal in log scale of Med15 binding in the indicated backgrounds. Targets increased and decreased when the C-terminus is deleted are indicated by color as in Fig. 2C. The color intensity corresponds to Pearson correlation to the binding of the full Med15 on the tested targets. (D-E) *Deleting Med2/3 together with the C-terminus of Med15 has no additional effect*. (D) Median signal of indicated backgrounds and mutations on the targets increased and decreased when the C-terminus is deleted (in log scale), The color intensity corresponds to Pearson correlation to Med15 ΔC400 on the represented targets. (E) Scatter plot showing the comparison of the binding signal across all promoters in the indicated strains as in Fig. 2A.

Alternatively, the C-terminus could direct Med15 binding through its interactions with other tail subunits. Consistent with this hypothesis, structural analyses place Med2, Med3, and Med5 in direct contact with the Med15 C-terminus. To test this possibility, we analyzed individual deletions and found that while *med5Δ* had no effect on Med15 binding, deletion of either *MED2* or *MED3* substantially reduced Med15 occupancy across all its target sites (Fig. 3B). Notably, the residual Med15 binding in both *med2Δ* and *med3Δ* backgrounds still localized to TF-enriched UASs upstream of the TSS and displayed a clear preference for the targets gained upon C-terminal truncation (Fig. 3C–E). Deletion of *MED16* further exacerbated the binding defects observed in both the *med3Δ* background and the C-terminally truncated mutant, further arguing against core Mediator coupling as the underlying mechanism (Fig. 3C). Furthermore, co-deletion of *MED2* and *MED3* in cells carrying the Med15 C-terminal mutation demonstrated epistasis, yielding no additional additive effects (Fig. 3D, E, Fig. S2B). Together, these results suggest that the C-terminus biases Med15 binding toward a subset of UASs favored by Med2 and Med3.

### The C-terminus can stabilize Med15 binding to TF-depleted targets

The three ABDs within the Med15 sequence direct genomic targeting through interactions with UAS-bound TFs. To examine whether the C-terminus similarly depends on these recruiting TFs, we utilized a genetic system that stabilizes Med15 binding at regions depleted of major recruiting TFs^26,32^. Specifically, we employed a strain lacking twelve key TFs normally present at Med15-bound targets, including the stress regulators Msn2 and Msn4 who’s binding closely correlated with Med15 occupancy under our experimental conditions (Fig. 4A, S2C). As expected, this multiplex TF deletion substantially reduced Med15 binding at target sites (Fig. 4B). We then sought to restore Med15 binding by fusing the protein to the heterologous DNA-binding domain (DBD) of Msn2 (Fig. 4A). Notably, while the Msn2-DBD alone exhibited limited occupancy at Med15 targets, the Med15-DBD fusion robustly localized to a large fraction of these TF-depleted UASs (Fig. S2C, Fig. 4C). Thus, in this engineered system, the DBD stabilizes overall Med15 genomic tethering, while the targeting specificity is conferred by Med15 itself.

**Figure 4:**
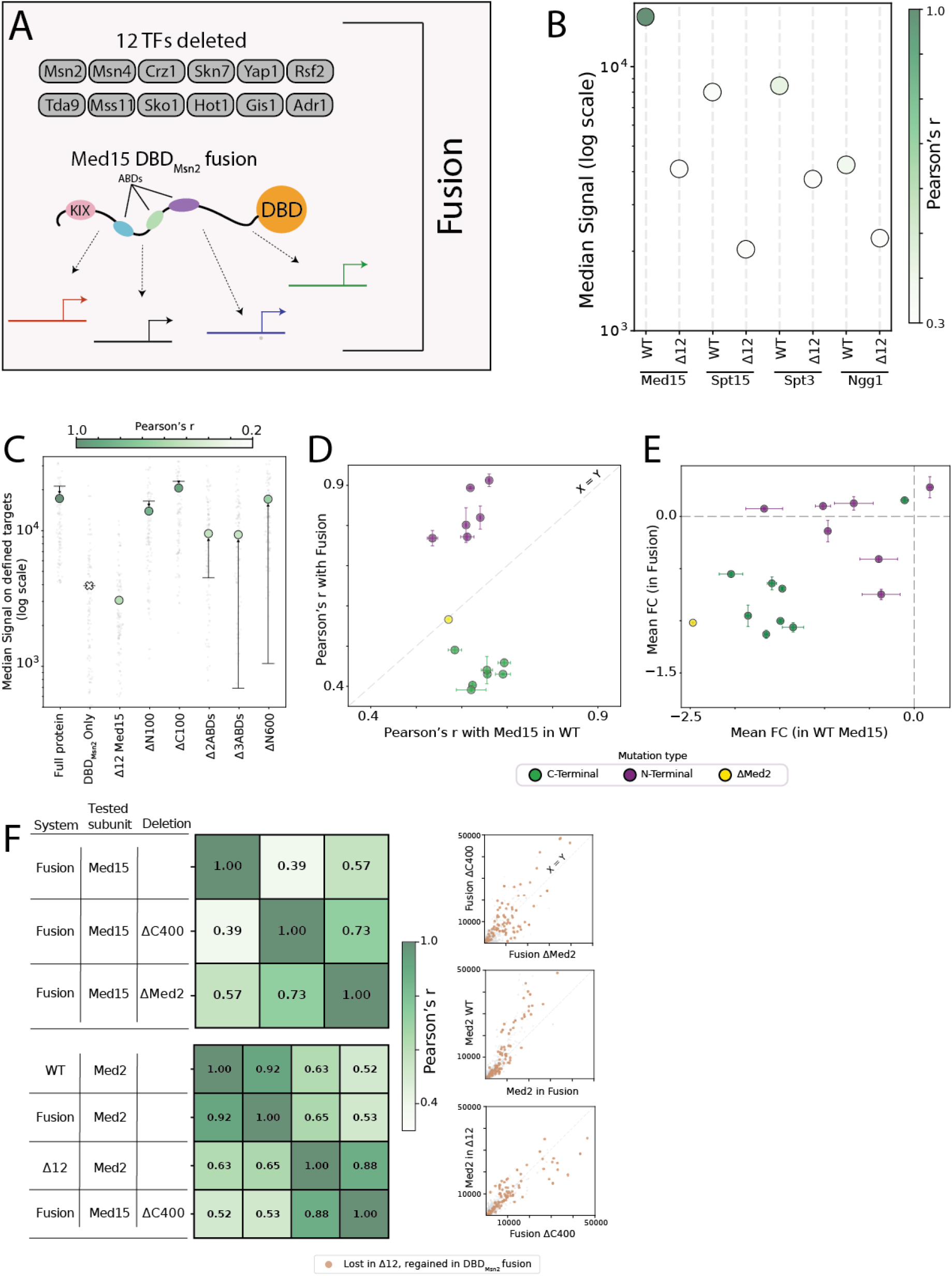
Mapping TF-dependent and independent elements contribute to Med15 binding preferences. (A) *The “Fusion” system: 12 TFs co-deleted system to study TF recruitment of cofactors with the addition of Med15 fused to DBD*_*Msn2*_. We used the previously described system in which the 12 indicated TFs were co-deleted, rendering a fraction of regulatory elements stripped of TFs to analyze cofactor recruitment. In the system called “fusion” we also added a fusion of DBD_Msn2_ to the C-terminus of Med15. (B) *Med15 and general cofactors do not localize to TF-depleted targets*. Median binding (in log scale) of Med15 and general cofactors in WT or TF-depleted cells. Color represents Pearson’s correlation to WT Med15. (C-E) *In targets lost in TF-depleted cells, but restored when Med15 is fused to DBD*_*Msn2*,_ *N-terminal mutations are rescued while C-terminal mutations do not*. (C) Median signals on targets lost in TF-depleted targets but regained in DBD-fusion (grey scattered dots) are presented in colored dots. The color represents the Pearson’s r to the full fused protein in the fusion system. The black horizontal lines represent the median signal of the WT Med15 protein containing the same indicated mutation on Med15 WT top targets. The black arrows indicate the changes in binding from the WT system to the fusion system. (D) A comparison of the correlation of the strains to the full Med15 in the two systems: fusion and WT is depicted. The diagonal gray line indicates where x = y. Green points represent C-terminal mutations, purple points N-terminal mutations and the yellow point represents the deletion of Med2. Error bars represent the STD. (E) Scatter plot comparing the median fold change (in log scale) from the full protein in the WT system to the same strain’s median fold change in the fusion system. Representation as in (D). (F) *Med2 deletion is similar to the C-terminal deletion while Med2 binding is retained in a TF-depleted system*. (Left) Shown are Pearson’s correlation coefficients comparing the binding signal of corresponding strains by system and tested subunit. (Right) Promoter binding signals of the indicated strains across all yeast promoters are shown. The diagonal dashed line indicates x=y. The brown dots represent the targets lost in the TF-depleted system and gained in the fusion, as shown in (C). (G)

To distinguish between TF-dependent and TF-independent recruitment, we focused on a subset of Med15 targets whose binding was abolished upon TF deletion but successfully rescued by the DBD fusion. We reasoned that Med15-DBD occupancy at these sites would become insensitive to mutations that impair natural TF recruitment, while remaining sensitive to perturbations affecting TF-independent binding mechanisms. Testing the impacts of N-terminal and C-terminal mutations within this background revealed distinct phenotypes. The defects caused by N-terminal mutations were largely compensated for in the TF-depleted context; notably, this included restoration of binding for a mutant entirely lacking the 600-residue N-terminal region that encompasses all three ABDs (Fig. 4C–E). By contrast, C-terminal mutations exhibited little, if any, compensation, resulting in a substantial shift in binding specificity across the TF-depleted UASs (Fig. 4C–E).

Given the similarity between the phenotypes of the C-terminal mutants and the med2Δ or med3Δ strains, we next asked whether Med2 and Med3 also influence Med15 binding across TF-depleted targets. Deletion of MED2 strongly impaired Med15–DBD occupancy at these sites. Moreover, the med2Δ mutation largely phenocopied the Med15 C-terminal truncations in both binding pattern and magnitude. Notably, this quantitative defect was moderate when compared to the near-complete loss of Med15 binding observed in wild-type cells under TF-depleted conditions (Fig. 4D–G).

These results support a model wherein the Med2-Med3 subcomplex acts through the Med15 C-terminus to direct genomic binding independently of major UAS-localized TFs. Consistent with this model, Med2 occupancy is retained (and in certain instances elevated) at TF-depleted UASs (Fig. 4F, G). Together, these findings indicate that Med2 and Med3 promote Med15 recruitment through both TF-dependent and TF-independent pathways, with the latter mechanism mediated, at least in part, by molecular interactions involving the Med15 C-terminus.

## Discussion

In this study, we investigated the molecular mechanisms underlying the genomic targeting of Med15. A central component of this process is recruitment by UAS-bound transcription factors (TFs), mediated by three activator-binding domains (ABDs) within Med15 that contact TF activation domains (ADs) through relatively weak interactions^14,15,17–19^. Because cooperation among weak interactions can enhance overall binding affinity, we hypothesized that Med15 targeting would require cooperative interactions among all three ABDs, rendering each domain individually essential. Contrary to this expectation, individual ABD deletions exerted only minor effects, and only their concurrent deletion completely abolished Med15 recruitment. Similarly, the disordered regions flanking the ABDs contributed modestly and redundantly. Together, these findings indicate that TF-Med15 interactions mediated by the ABDs remain weak and are not organized into a strongly cooperative system. Consequently, these ABDs alone are insufficient to account for the genome-wide recruitment pattern of Med15.

Instead, we identified an intrinsically disordered region within the Med15 C-terminus that plays a pivotal role in shaping Med15 binding across UASs. Removal of this region redistributed Med15 occupancy, reducing binding at a large subset of targets while increasing it at others, despite the presence of intact ABDs. Because this region overlaps with the Mediator-association domain (MAD) required for Mediator integrity^28,29,7^, we initially hypothesized that it biases Med15 recruitment through interactions with the core Mediator complex, which associates broadly with promoters independently of the TF-recruited tail module. However, this model was refuted by *MED16* deletion; while Med16 decouples the tail module from the core Mediator, its absence caused only minor effects on Med15 occupancy and failed to recapitulate the C-terminal truncation phenotype.

Our results instead point to Med2 and Med3 as the functional partners of the Med15 C-terminus. These subunits assemble into an independent Med2-Med3-Med15 tail triad^31,33^, and multiple lines of evidence indicate that they underlie the C-terminal contribution to UAS targeting. First, deletion of *MED2* or *MED3* strongly reduced Med15 occupancy, while partially phenocopying the redistribution pattern observed upon C-terminal truncation. Second, combining *MED2* or *MED3* deletion with the C-terminal truncation did not exacerbate the targeting defect, consistent with a shared epistatic pathway. Third, under TF-depleted conditions, *med2Δ* cells closely phenocopied the effects of Med15 C-terminal mutations on genomic binding patterns. Together, these findings support a model wherein Med2–Med3 interactions with the Med15 C-terminus bias Med15 binding toward a specific subset of UASs.

Our TF-depletion and rescue system further revealed that the Med15 C-terminal region, together with Med2 and Med3, continues to shape Med15 binding even at UASs depleted of major recruiting TFs. In this engineered system, Med15 occupancy could be restored by heterologous DBD tethering, yet binding preferences remained strictly dependent on the C-terminal region and the Med2–Med3 subcomplex, whereas the ABDs exerted only minor effects. Consistently, Med2 remains bound at TF-depleted UASs that have lost Med15 occupancy. This baseline binding could reflect recruitment by additional TFs not included in our multiplex deletion strain, although we targeted the twelve TFs showing the highest correlation with wild-type Med15 binding. Alternatively, Med2 or the Med15 C-terminus may recognize these UAS sequences independently of TFs, perhaps through IDR-directed interactions^34–36^. Notably, the specific UASs affected by C-terminal truncation (or *MED2* deletion) differ between wild-type and TF-depleted conditions, likely reflecting differences in structural stabilization provided by the fused DBD versus natural, UAS-bound TFs.

An unexpected observation from our study is that the C-terminal truncation partially compensates for the loss of Med2 or Med3, resulting in less severe reductions in overall binding compared to *MED2* or *MED3* deletions alone. This compensatory effect is unlikely to arise from altered coupling to the core Mediator, as *MED16* deletion exacerbates rather than rescues these deletion phenotypes. One possibility is that, in the absence of its C-terminal region, Med15 adopts an alternative interaction mode that partially bypasses the requirement for Med2 and Med3. Alternatively, the C-terminal region may impose structural or autoinhibitory constraints on Med15 binding that are relieved upon truncation. Elucidating the precise mechanistic basis of this compensation will require further structural and biophysical investigation.

Together, our findings support a model in which Med15 recruitment is governed by two mechanistically distinct layers (Fig. 5): weak, redundant ABD-mediated interactions that enable initial TF-dependent recruitment, and C-terminal interactions with Med2–Med3 that shape genomic binding specificity independently of TFs. This division of labor provides a conceptual framework for understanding how the Mediator complex integrates diverse TF signals while maintaining highly selective genome targeting. More broadly, our results highlight the critical importance of non-ABD regions in Mediator function and reveal an unappreciated layer of regulation in transcriptional coactivator recruitment *in vivo*.

**Figure 5:**
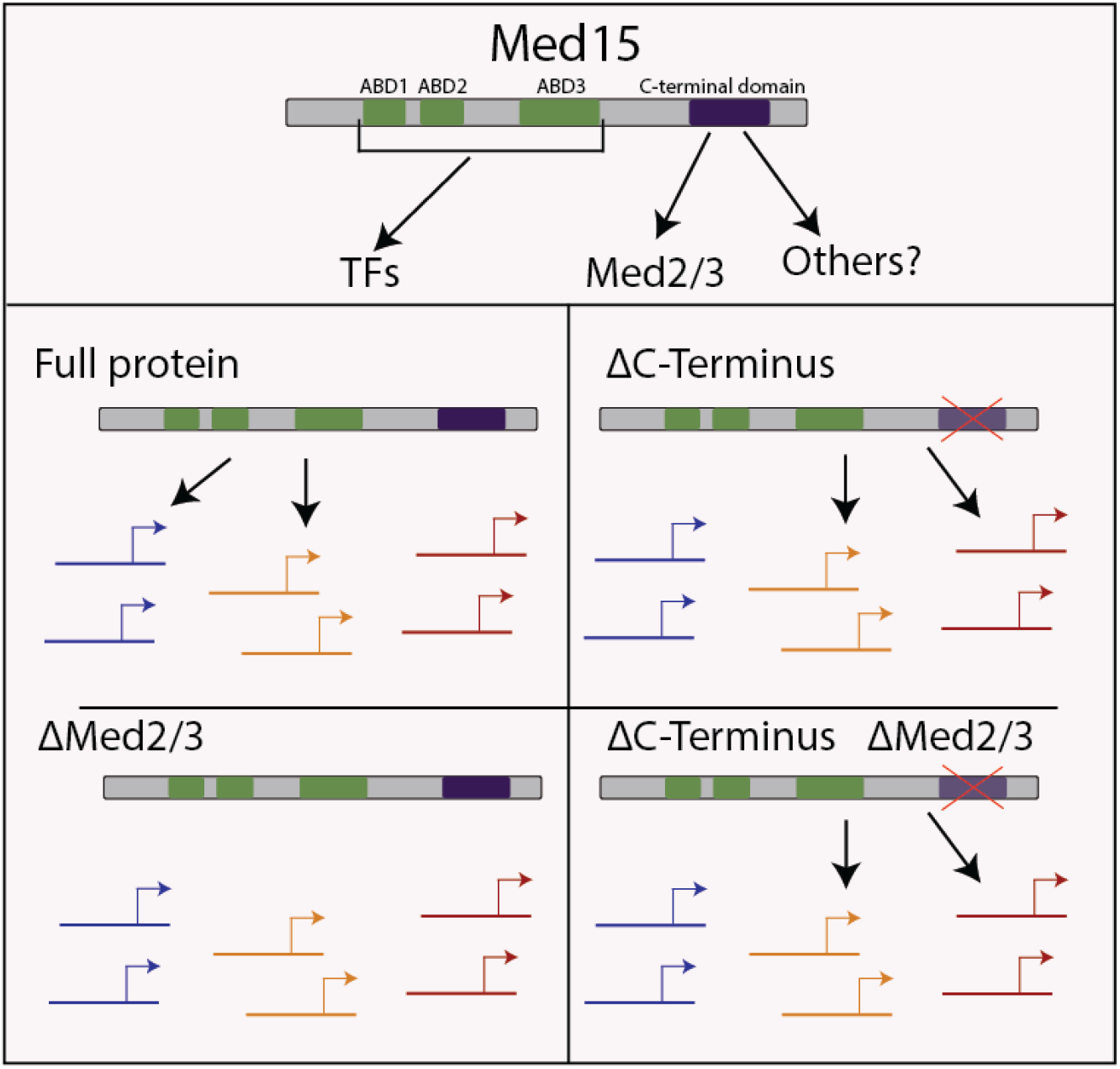
*Proposed summary scheme* (see Discussion). Presented are the sequence determinants of Med15 and the tail subcomplex that influence the binding preferences of Med15 and the possible dependencies each sequence element has.

## Acknowledgments

We want to thank past and current members of the Barkai lab for their assistance with this project. We specifically want to thank Dr. Sagie Brodsky and Dr. Offir Lupo for the technical help, the wonderful comments, and all of the support throughout this project.

This project was supported by the European Research Council (ERC), Israel Science Foundation (ISF), German Research Foundation (DFG), and the Minerva Center. The project is also supported by the Adams Fellowships Program of the Israel Academy of Sciences and Humanities. A.C. is a fellow of the Ariane de Rothschild Women Doctoral Program.

## Author contributions

Conceptualization: A.C., V.M., N.B.; Methodology: A.C., G.Y., N.B.; Investigation: A.C., J.B.; Visualization and analysis: A.C., N.B.; Funding acquisition: N.B.; Supervision: N.B.; Writing – original draft: A.C., N.B.; Writing – review & editing: A.C., V.M., G.Y., N.B.

## Data availability

The link to the code and notebooks to produce all of the figures: https://doi.org/10.5281/zenodo.16641455

Generated high-throughput sequencing data is available in the GEO under the GSE292501 accession number.

---

**Figure S1.**
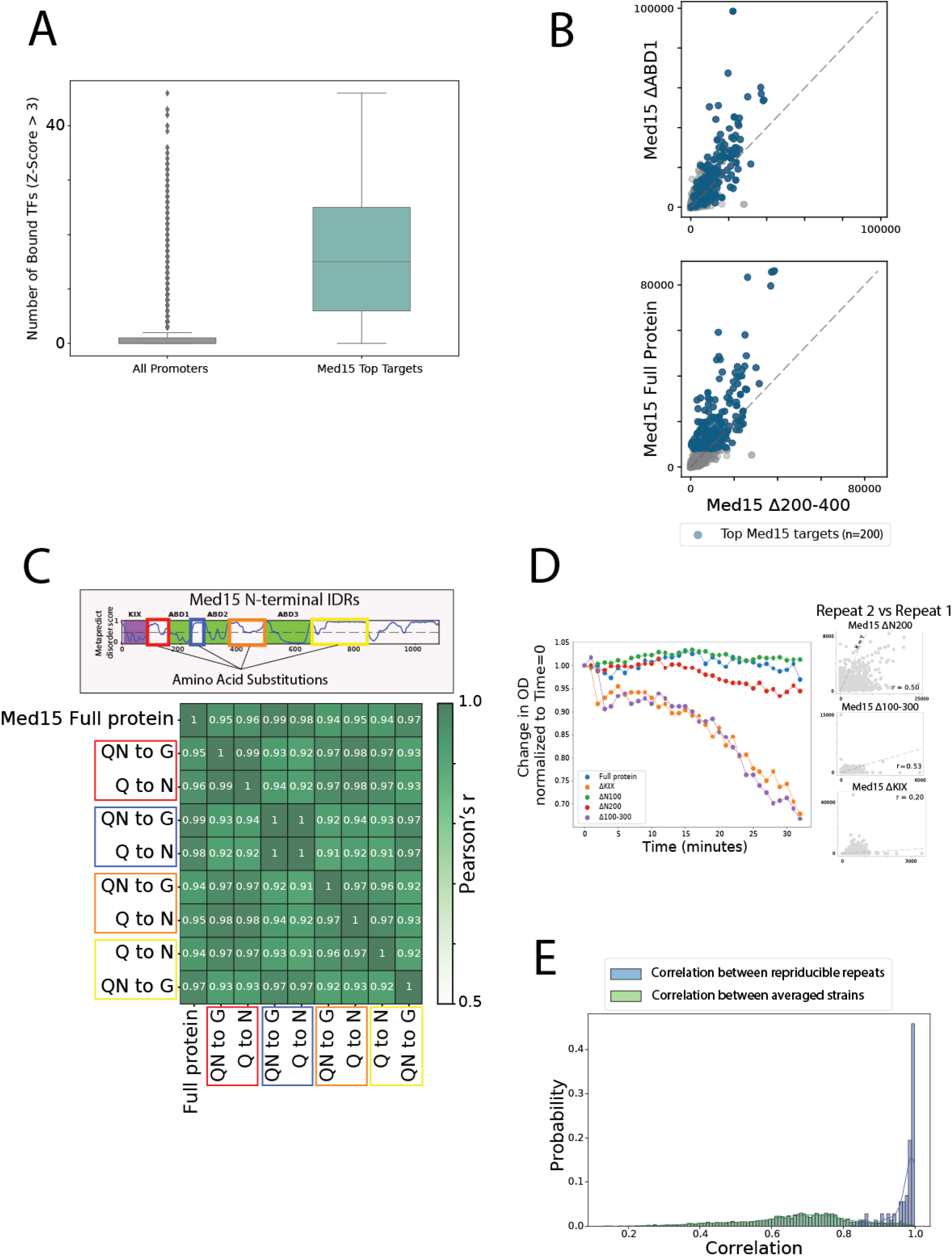
(A) *Med15 targets are TF-dense*. Boxplot comparing the amount of TFs bound to top Med15 targets versus all promoters in the genome. This was tested by calculating the Z score of each TF on the promoters and comparing the ones above 3 between the target groups. (A) *The deletion of ABD1 and ABD2 (the deletion of segment 200-400AA) is similar to the deletion of ABD1 alone*. Binding of indicated strains across all promoter is depicted with a scatter plot. The diagonal dashed line indicates x=y. Blue dots represent the top Med15 targets shown in Fig. 1B. (B) *Amino acid substitutions in the IDRs in the N-terminus of Med15 don’t change binding preference*. Shown are Pearson’s r correlation coefficients comparing the binding signal of the indicated strains. (top) Depicted are the N-terminal disordered regions that were mutated, each represented by a different corresponding color. (C) *Non-reproducible repeats exhibit a flocculation phenotype*. (left) a flocculation assay was performed on the Med15 strains studied in this paper (methods) and the change in OD was measured for 33 time points. The OD was normalized according to time=0 minutes and plotted, the flocculant strains are compared to the full Med15 and to Med15 ΔN100 which doesn’t show a flocculation phenotype. (right) Representative scatter plots of the non-reproducible repeats are shown, with repeat 1 being plotted on the x-axis and repeat 2 plotted on the y-axis. The diagonal dashed line indicates the x=y line. (D) *Reproducible repeats present a high correlation between them, while between the strains themselves, the correlation coefficients vary*. Pearson’s r correlation coefficient was calculated between experimental repeats (blue), coefficients above 0.8 were used, and the repeats were averaged. The correlation coefficient between averaged repeats (different strains) was calculated (green) and a variation is shown

**Figure S2.**
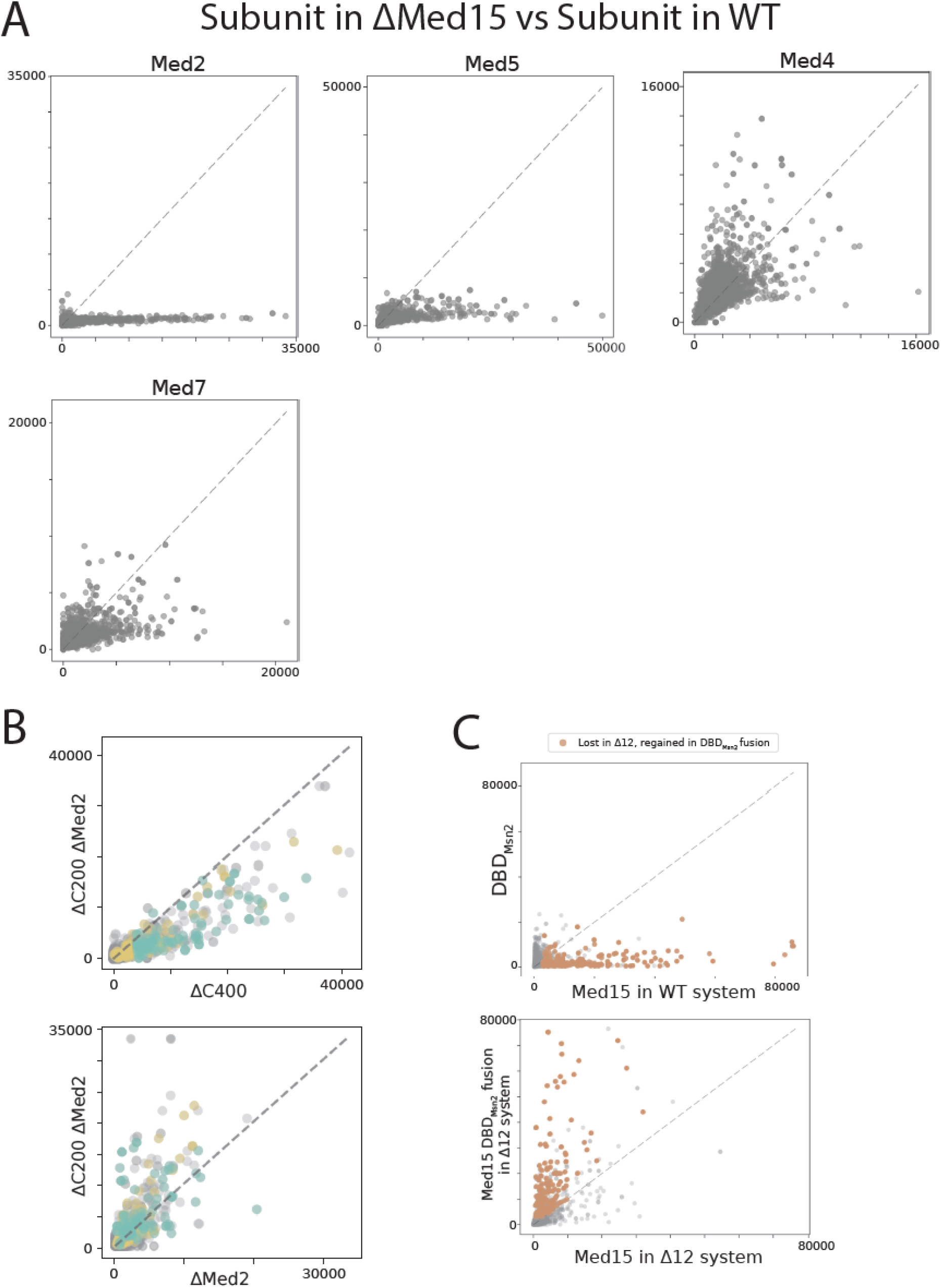
(A) *Med15 affects the genomic localization of tail and middle subunits of the mediator complex*. Presented are scatter plots comparing tail subunits and middle subunits in WT cells (x-axis) and when Med15 is deleted (y-axis). Representation as in Fig. S1B. (B) *Deleting Med2/3 together with the C-terminus of Med15 has no additional effect*. (top) Median signal of indicated backgrounds and mutations on the targets increased and decreased when the C-terminus is deleted (in log scale), The color intensity corresponds to Pearson correlation to Med15 ΔC400 on the represented targets. (bottom) Scatter plot showing the comparison of the binding signal across all promoters in the indicated strains as in Fig. 2A. **(C)** *DBD*_*Msn2*_ *binding differs from Med15 binding in the WT system, but the fusion of DBD*_*Msn2*_ *to Med15 rescues binding in TF depleted targets*. Presented are scatter plots depicting the binding signal across all yeast promoters of the indicated strains. Presentation as in Fig. 4F.

**Figure S3.**
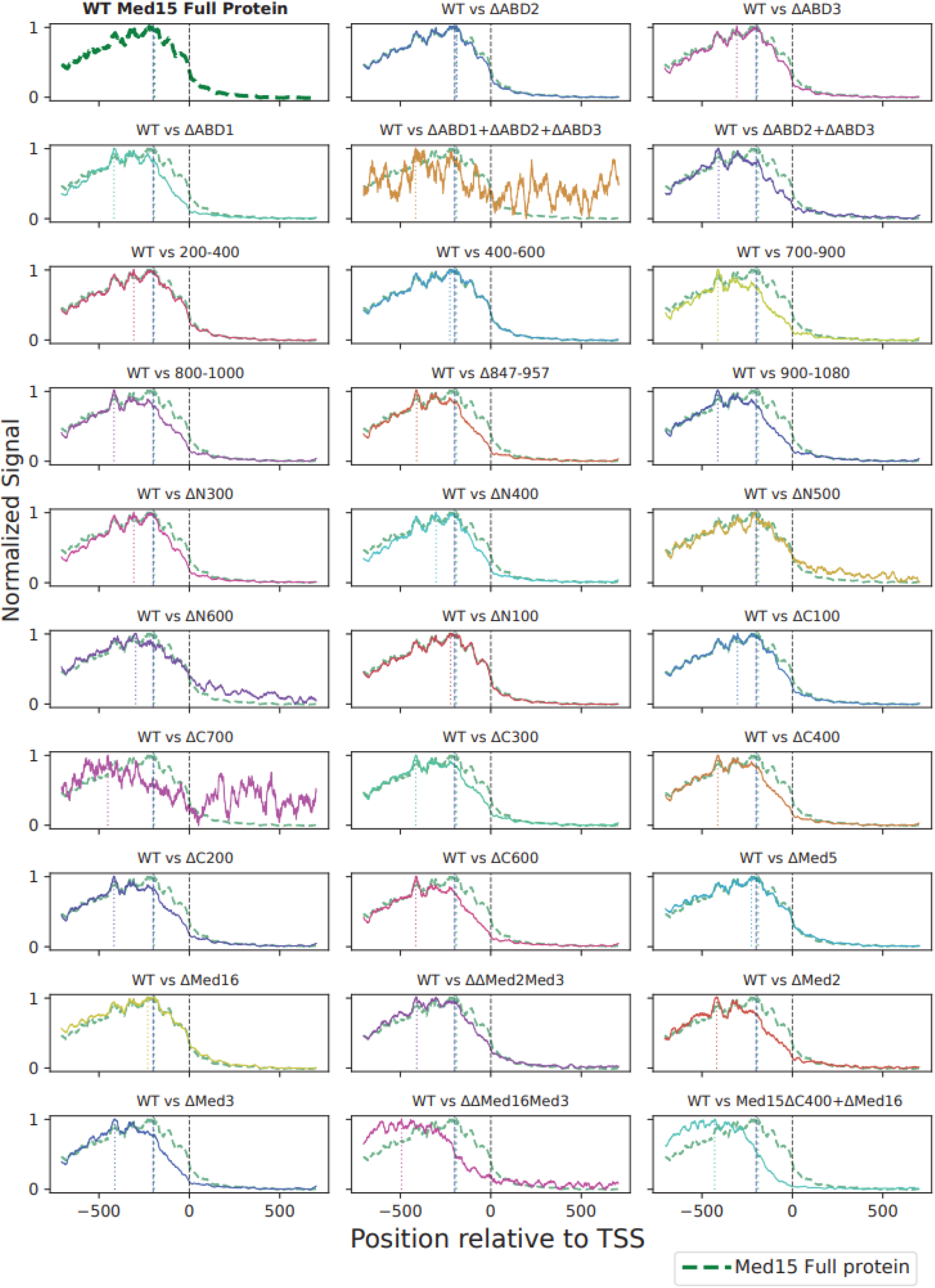
Binding signals of the indicated strains compared to Med15 WT (green, dashed) at the top Med15 targets 200 targets are shown as meta profiles centered around the TSS and flanked by 700bp regions upstream and downstream (methods, Table S1).

